## Supplemental material for "Structural and molecular biology of Acheta domesticus segmented densovirus, the first parvovirus to harbor a bipartite genome"

**S1 Table:** The exact positions of the open reading frames (ORFs) located on the NS and VP segments of Acheta domesticus segmented densovirus, respectively.

| **NS segment** | | |
| --- | --- | --- |
| **ORF name** | **Start position** | **End position** |
| ORF1 (NS1) | 471 nt | 2858 nt |
| ORF2 (NS2) | 559 nt | 1695 nt |
| ORF3 | 1756 nt | 1824 nt |
| ORF4 (NS1-N1 2^nd^ exon) ^a^ | 2177 nt | 2209 nt |
| ORF5 (NS1-N2 2^nd^ exon) ^a^ | 2351 nt | 2500 nt |

^a^ORF2 has no ATG start codon

| **VP segment** | | |
| --- | --- | --- |
| **ORF name** | **Start position** | **End position** |
| ORF1 | 483 nt | 1562 nt |
| ORF2 | 1543 nt | 1854 nt |
| ORF3 | 1859 nt | 2899 nt |

**S2Table:** Intron and exon boundaries of the Acheta domestica segmented densovirus spliced transcripts

| **Transcript exon** | **Start pos.** | **End pos.** | **Transcript intron** | **Start pos.** | **End pos.** |
| --- | --- | --- | --- | --- | --- |
| **NS segment** | | | | | |
| Transcript 2, exon 1 | 579 nt | 1391 nt | Transcript 2, intron 1 | 1392 nt | 2185 nt |
| Transcript 2, exon 2 | 2186 nt | 2209 nt |  |  |  |
| Transcript3, exon 1 | 579 nt | 1818 nt | Transcript 3, intron 1 | 1819 nt | 2423 nt |
| Transcript 3, exon 2 | 2424 nt | 2500 nt |  |  |  |
| Transcript5, exon 1 | 949 nt | 1391 nt | Transcript 5, intron 1 | 1392 nt | 2185 nt |
| Transcript 5, exon 2 | 2186 nt | 2858 nt |  |  |  |
| Transcript6, exon 1 | 1756 nt | 1818 nt | Transcript 6, intron 1 | 1819 nt | 2423 nt |
| Transcript 6, exon 2 | 2424 nt | 2858 nt |  |  |  |
| **VP segment** | | | | | |
| Transcript 3, exon 1 | 1543 nt | 1752 nt | Transcript 3, intron 1 | 1753 nt | 2032 nt |
| Transcript 3, exon 2 | 2033 nt | 2899 nt |  |  |  |
| Transcript 4, exon 1 | 1543 nt | 1752 nt | Transcript 4, intron 1 | 1753 nt | 2611 nt |
| Transcript 4, exon 2 | 2612 nt | 2899 nt |  |  |  |

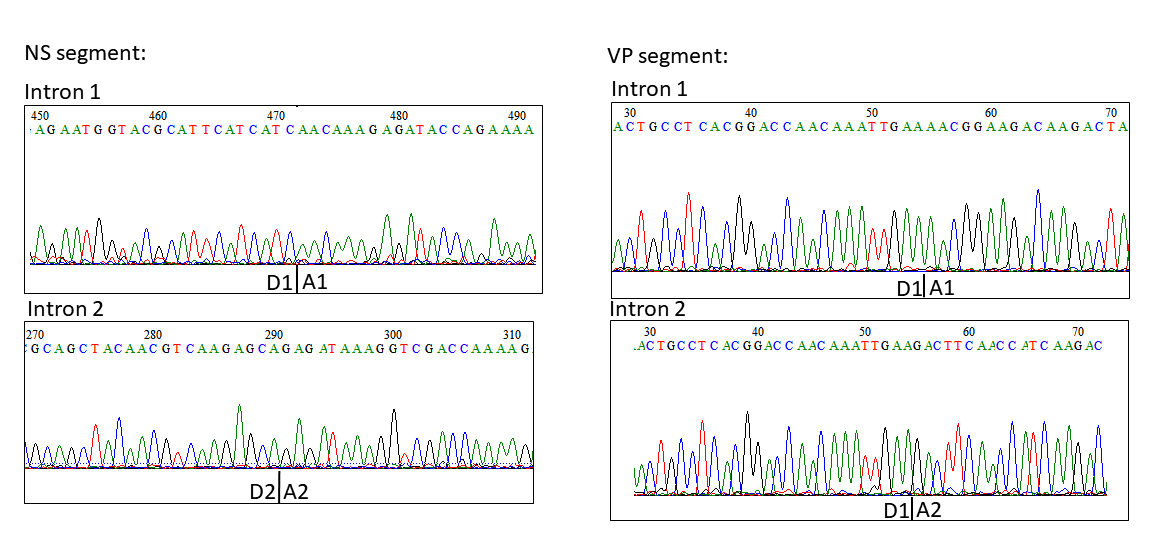

**S3 Figure:** Electrophoretograms showing the spliced-out introns of sequenced cDNA of mRNA derived from AdSDV-infected common house crickets.

**S4 Table:** Protein sequencing results of the nano-liquid chromatography tandem mass spectrometry, searched against the complete translated Acheta domesticus segmented densovirus (AdSDV) genome as well as the NCBI non-redundant protein sequence database.

55 kDa, HB band

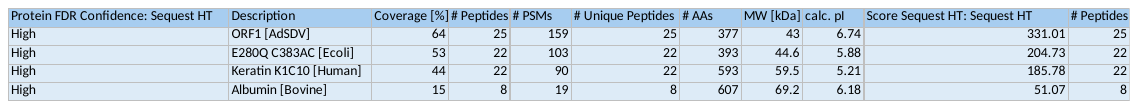

50 kDa, HB band

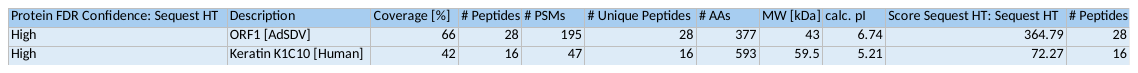

43 kDa, HB band

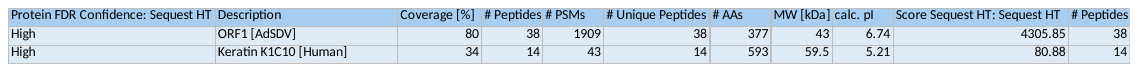

55 kDa, LB band

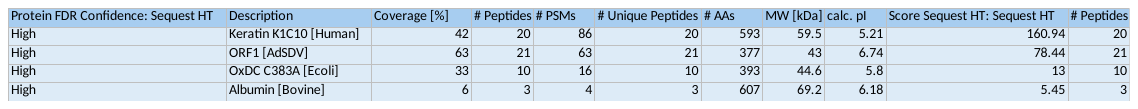

50 kDa, LB band

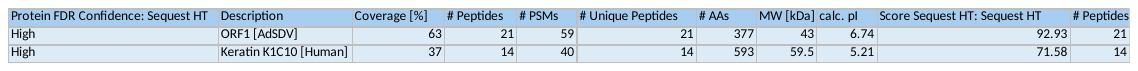

43 kDa, LB band

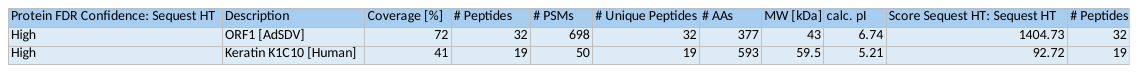

38 kDa, LB band

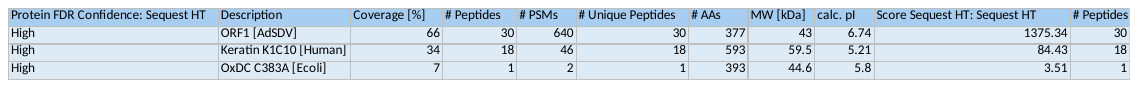

Normalized RFU

| Elapsed time | ORF1 | HB | LB | B19 VLPs | ORF1 heated | HB heated | LB heated | negative control |
| --- | --- | --- | --- | --- | --- | --- | --- | --- |
| 0:00:00 | 1 | 1 | 1 | 1 | 1 | 1 | 1 | 1 |
| 0:01:00 | 0.988636 | 0.989691 | 1.0102 | 1 | 0.990099 | 0.973451 | 0.971154 | 1.01786 |
| 0:02:00 | 1 | 1 | 1.0102 | 1.0099 | 0.980198 | 0.964602 | 0.971154 | 1.01786 |
| 0:03:00 | 1 | 1 | 1.0102 | 1.0198 | 0.980198 | 0.964602 | 1 | 1.01786 |
| 0:04:00 | 1 | 1 | 0.989796 | 1.0396 | 0.980198 | 0.973451 | 1 | 1.01786 |
| 0:05:00 | 1 | 1.01031 | 0.979592 | 1.0495 | 0.970297 | 0.973451 | 0.990385 | 1.01786 |
| 0:06:00 | 1 | 1.01031 | 0.969388 | 1.05941 | 0.980198 | 0.973451 | 0.990385 | 1.01786 |
| 0:07:00 | 1 | 1.01031 | 0.969388 | 1.07921 | 0.970297 | 0.982301 | 0.990385 | 1.01786 |
| 0:08:00 | 1 | 1.02062 | 0.969388 | 1.08911 | 0.980198 | 0.982301 | 0.990385 | 1 |
| 0:09:00 | 1 | 1.01031 | 0.969388 | 1.09901 | 0.980198 | 0.982301 | 0.990385 | 1.01786 |
| 0:10:00 | 1 | 1 | 0.969388 | 1.11881 | 0.980198 | 0.99115 | 0.990385 | 1.01786 |
| 0:11:00 | 1 | 0.989691 | 0.969388 | 1.12871 | 0.990099 | 0.99115 | 1 | 1.01786 |
| 0:12:00 | 1 | 0.989691 | 0.979592 | 1.14851 | 1 | 0.99115 | 1 | 1.01786 |
| 0:13:00 | 1 | 0.989691 | 0.979592 | 1.15842 | 1 | 0.982301 | 1.00962 | 1.01786 |
| 0:14:00 | 1 | 0.989691 | 0.989796 | 1.16832 | 0.990099 | 0.982301 | 1.00962 | 1.01786 |
| 0:15:00 | 1 | 1 | 0.989796 | 1.18812 | 0.990099 | 0.99115 | 1.00962 | 1.01786 |
| 0:16:00 | 1 | 1.01031 | 0.989796 | 1.20792 | 0.980198 | 0.982301 | 1.00962 | 1 |
| 0:17:00 | 1 | 1.01031 | 0.989796 | 1.21782 | 0.980198 | 0.982301 | 1.00962 | 1.01786 |
| 0:18:00 | 1 | 1.02062 | 0.989796 | 1.22772 | 0.980198 | 0.982301 | 1.00962 | 1.01786 |
| 0:19:00 | 1 | 1.02062 | 0.989796 | 1.23762 | 0.980198 | 0.982301 | 1 | 1.01786 |
| 0:20:00 | 1 | 1.01031 | 0.979592 | 1.23762 | 0.980198 | 0.982301 | 1 | 1.01786 |
| 0:21:00 | 0.988636 | 1.01031 | 0.979592 | 1.24752 | 0.980198 | 0.99115 | 0.990385 | 1.01786 |
| 0:22:00 | 1 | 1 | 0.979592 | 1.24752 | 0.980198 | 0.982301 | 0.980769 | 1 |
| 0:23:00 | 1 | 1 | 0.979592 | 1.25743 | 0.980198 | 0.982301 | 0.980769 | 1.01786 |
| 0:24:00 | 1 | 1 | 0.979592 | 1.25743 | 0.980198 | 0.99115 | 0.980769 | 1.01786 |
| 0:25:00 | 0.988636 | 1 | 0.979592 | 1.26733 | 0.980198 | 0.99115 | 0.980769 | 1 |
| 0:26:00 | 1 | 1 | 0.979592 | 1.27723 | 0.980198 | 0.99115 | 0.980769 | 1.01786 |
| 0:27:00 | 1 | 1 | 0.979592 | 1.28713 | 0.980198 | 0.99115 | 0.980769 | 1 |
| 0:28:00 | 1 | 1.01031 | 0.979592 | 1.29703 | 0.980198 | 1 | 0.980769 | 1.01786 |
| 0:29:00 | 1 | 1.01031 | 0.979592 | 1.31683 | 0.980198 | 1 | 0.980769 | 1.01786 |
| 0:30:00 | 1 | 1.01031 | 0.979592 | 1.32673 | 0.980198 | 1 | 0.980769 | 1 |
| 0:31:00 | 1 | 1.01031 | 0.989796 | 1.34653 | 0.990099 | 1 | 0.980769 | 1.01786 |
| 0:32:00 | 1 | 1.01031 | 0.989796 | 1.35644 | 0.990099 | 1 | 0.980769 | 1 |
| 0:33:00 | 1 | 1.01031 | 0.989796 | 1.36634 | 0.990099 | 1 | 0.980769 | 1.01786 |
| 0:34:00 | 1 | 1.02062 | 0.989796 | 1.37624 | 0.990099 | 1 | 0.980769 | 1.01786 |
| 0:35:00 | 1 | 1.01031 | 0.989796 | 1.38614 | 0.990099 | 1 | 0.980769 | 1.01786 |
| 0:36:00 | 1.01136 | 1.02062 | 0.989796 | 1.39604 | 0.990099 | 1 | 0.980769 | 1.01786 |
| 0:37:00 | 1.01136 | 1.02062 | 0.989796 | 1.40594 | 0.990099 | 1 | 0.980769 | 1.01786 |
| 0:38:00 | 1.01136 | 1.02062 | 0.989796 | 1.41584 | 0.990099 | 1 | 0.980769 | 1.01786 |
| 0:39:00 | 1.01136 | 1.01031 | 1 | 1.43564 | 0.990099 | 1 | 0.980769 | 1.01786 |
| 0:40:00 | 1.01136 | 1.02062 | 1 | 1.44554 | 0.990099 | 1 | 0.980769 | 1.01786 |
| 0:41:00 | 1.01136 | 1.01031 | 1 | 1.45545 | 0.990099 | 1 | 0.980769 | 1.01786 |
| 0:42:00 | 1.01136 | 1.01031 | 1 | 1.46535 | 0.990099 | 1.00885 | 0.980769 | 1.01786 |
| 0:43:00 | 1.01136 | 1.02062 | 1 | 1.47525 | 0.990099 | 1.00885 | 0.980769 | 1.01786 |
| 0:44:00 | 1.01136 | 1.01031 | 1 | 1.48515 | 0.990099 | 1.00885 | 0.980769 | 1.01786 |
| 0:45:00 | 1.01136 | 1.02062 | 1 | 1.49505 | 0.990099 | 1.00885 | 0.980769 | 1 |
| 0:46:00 | 1.01136 | 1.02062 | 1 | 1.50495 | 0.990099 | 1.00885 | 0.980769 | 1.01786 |
| 0:47:00 | 1.01136 | 1.02062 | 1 | 1.51485 | 0.990099 | 1.00885 | 0.980769 | 1.01786 |
| 0:48:00 | 1.01136 | 1.02062 | 1 | 1.52475 | 0.990099 | 1.00885 | 0.980769 | 1.01786 |
| 0:49:00 | 1.01136 | 1.02062 | 1 | 1.53465 | 0.990099 | 1.00885 | 0.980769 | 1 |
| 0:50:00 | 1.01136 | 1.02062 | 1 | 1.54455 | 0.990099 | 1.00885 | 0.980769 | 1.01786 |
| 0:51:00 | 1.01136 | 1.02062 | 1 | 1.55446 | 0.990099 | 1.00885 | 0.980769 | 1.01786 |
| 0:52:00 | 1.01136 | 1.02062 | 1 | 1.56436 | 0.990099 | 1.00885 | 0.980769 | 1.01786 |
| 0:53:00 | 1.01136 | 1.02062 | 1 | 1.57426 | 0.990099 | 1.00885 | 0.980769 | 1.01786 |
| 0:54:00 | 1.01136 | 1.02062 | 1 | 1.58416 | 0.990099 | 1.00885 | 0.980769 | 1.01786 |
| 0:55:00 | 1.01136 | 1.02062 | 1.0102 | 1.60396 | 1 | 1.00885 | 0.980769 | 1.01786 |
| 0:56:00 | 1.01136 | 1.02062 | 1.0102 | 1.61386 | 1 | 1.0177 | 0.980769 | 1.01786 |
| 0:57:00 | 1.01136 | 1.03093 | 1.0102 | 1.62376 | 1 | 1.0177 | 0.980769 | 1.01786 |
| 0:58:00 | 1.01136 | 1.03093 | 1.0102 | 1.62376 | 0.990099 | 1.0177 | 0.980769 | 1.01786 |
| 0:59:00 | 1.01136 | 1.03093 | 1.0102 | 1.64356 | 1 | 1.0177 | 0.980769 | 1.01786 |
| 1:00:00 | 1.01136 | 1.03093 | 1.0102 | 1.65347 | 1 | 1.0177 | 0.980769 | 1.01786 |

**S5 Figure:** Colorimetric assay measuring the PLA2 activity of the Acheta domestica segmented densovirus (AdSDV) capsids, using parvovirus B19 virus like particles (VLPs) as positive control. Values are normalized to the ones measured at the first timepoint of the assay. HB abbreviates the high buoyancy capsids pulled from the 20% band of the sucrose step gradient, used for AdSDV purification. LB (low buoyancy) capsids derive from the 30% band of the aforementioned sucrose step gradient. ORF1 indicates VLPs assembled exclusively from the protein product of VP ORF1, lacking the PLA2 domain, and functions as negative control. Purified particles were heated to 60^◦^C to obtain the data indicated as “heated”.

**S6 Table:** Data collection and refinement statistics of the Acheta domestica segmented densovirus capsids and ORF1 only virus-like particles (VLPs)

| **Processing and Refinement Parameters** | LB | HB1 | HB2 | HB | ORF1 |
| --- | --- | --- | --- | --- | --- |
| Total number of micrographs | 1411 | 970 | 1002 | 1127 | 1700 |
| Reconstruction software | cisTEM | cisTEM | cisTEM | cisTEM | cisTEM |
| Defocus range (µm) | 0.65-4.00 | 0.81-4.00 | 0.8-4.32 | 0.62-4.41 | 0.57-2.57 |
| Electron dose (e^−^/Å^2^) | 75 | 60 | 60 | 75 | 60 |
| Frames/micrograph | 50 | 50 | 50 | 50 | 50 |
| Pixel size (Å/pixel) | 1.041 | 1.038 | 1.038 | 1.041 | 1.049 |
| Starting number of particles | 188074 | 46082 | 58680 | 98920 | 18598 |
| Particles used for final map | 150469 | 30065 | 48354 | 59211 | 13019 |
| B-factor used for final map (Å^2^) | 20 (Post-Cut-Off B-factor) | 10 (Post-Cut-Off B-factor) | 10 (Post-Cut-Off B-factor) | 20 (Post-Cut-Off B-factor) | 20 (Post-Cut-Off B-factor) |
| Resolution of final map (Å) | 19 | 3 | 3.1 | 2.5 | 3.3 |
| Residue range (VP1) | 47-377 | 49-366 | 49-366 | 49-366 | 49-366 |
| Map correlation coefficient | 0.6528 | 0.629 | 0.6115 | 0.7147 | 0.6774 |
| RMSD (root-mean-square deviation) [bonds] (Å) | 0.010 | 0.008 | 0.011 | 0.009 | 0.011 |
| RMSD [angles] (Å) | 0.852 | 0.788 | 0.863 | 0.772 | 0.849 |
| All-atom clash score | 12.19 | 9.35 | 12.7 | 15.3 | 13.6 |
| Favored (%) | 95 | 98.1 | 97.5 | 96.1 | 95.9 |
| Allowed (%) | 4.5 | 1.6 | 2.5 | 3.6 | 3.8 |
| Outliers (%) | 0.5 | 0.3 | 0 | 0.3 | 0.3 |
| Rotamer outliers (%) | 0 | 0.3 | 0 | 0 | 0 |
| C-β deviations | 0 | 0 | 0 | 0 | 0 |

|  | **Inner radius (Å)** | **Inner surface area (nm^2^)** | **Inner volume (nm^3^)** | **Genome size (nt)** | **Genus** |
| --- | --- | --- | --- | --- | --- |
| **CPV** | 92.9 | 1085 | 3360 | 5323 | *Protoparvovirus* |
| **AAV2** | 89.9 | 1015 | 3040 | 4679 | *Dependoparvovirus* |
| **GmDV** | 98.7 | 1,223 | 4,023 | 6039 | *Protoambidensovirus* |
| **AdDV** | 91.8 | 1,056 | 3,228 | 5425 | *Scindoambidensovirus* |
| **BmDV1** | 98.7 | 1224 | 4,028 | 5076 | *Iteradensovirus* |
| **PstDV** | 87.6 | 963 | 2,811 | 3914 | *Penstylhamaparvovirus* |
| **AdSDV** | 82.2 | 848 | 2322 | 3332 / 3316 | *Brevihamaparvovirus* |
| **PmMDV** | 86.5 | 933 | 2710 | 4371 | Incertiparvovirinae |

**A**

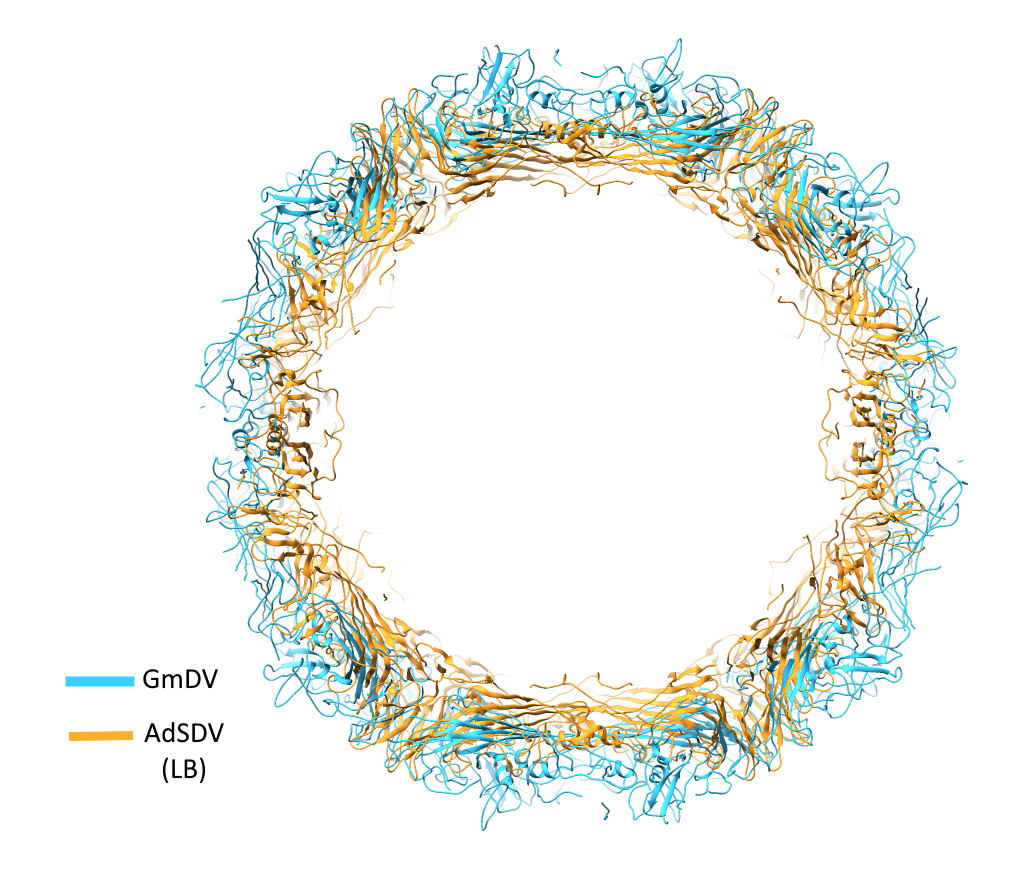

**B**

**S7 Figure:** Comparing the capsid and gnome metrics of various members of the Parvoviridae, with those of Acheta domesticus segmented densovirus (AdSDV). Subfamilies, including the Parvovirinae, Densovirinae and Hamaparvovirinae, respectively, are separated by the thick lines. Abbreviations: CPV – canine parvovirus, AAV – Adeno-associated virus 2, GmDV – Galleria mellonella densovirus, AdDV – Acheta domestica densovirus, BmDV1 – Bombyx mori densovirus 1, PdtDV – Penaeus stylirostris densovirus, PmMDV – Penaeus stylirostris metallodensovirus

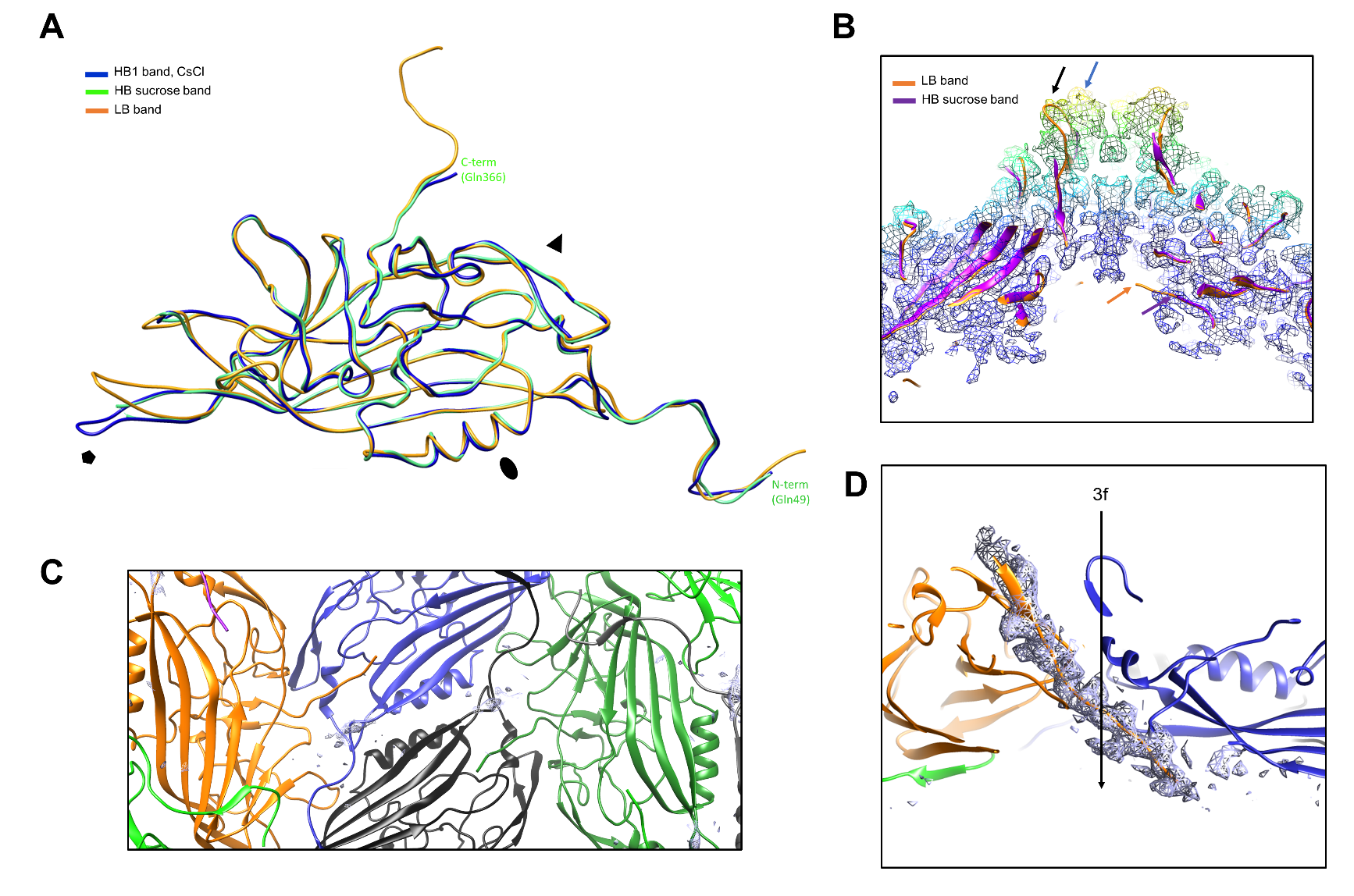

**S8 Figure:** The high buoyancy (HB) band, pulled from the sucrose step gradient, contains a mixed population on Acheta domestica segmented densovirus (AdSDV) capsids. (A) When superimposing the structural model of the HB band monomer with those of the HB bands pulled from the continuous CsCl gradient and with the low buoyancy (LB) capsid monomers, the HB sucrose capsids display the disordered N- and C termini of the CsCl HB capsids, accompanied by a disordered DE loop, as result of the conformation difference between the LB and HB1-2 populations. (B) Cross section of the HB sucrose capsid fivefold axis, colored radially, with the electron density indicated by the colored mash (ơ=1). The model built into the density is shown as ribbon diagrams, superimposed with that of the LB capsids. The HB sucrose capsids display the DE loop density characteristic for both the LB fivefold axis (black arrow) and the CsCl HB1 and 2 fivefold axes (blue arrow). Note the density-filled fivefold channel, which lacks the connection to the first N-terminally ordered residue (orange arrow for LB, magenta arrow for HB sucrose capsids), unlike in case of the LB capsids. (C) The ssDNA-binding luminal region, located directly under the twofold symmetry axis, shown as ribbon diagrams. Display of the electron density map was zoned to the nucleic acid models exclusively, indicating the lack of ordered nucleotides (ơ=1). (D) Side view of the HB sucrose capsid threefold annulus, with the black arrow indicating the location of the threefold axis. The density, displayed as the purple mash and zoned on the final C-terminal 20 residues, indicates that the ordered segment ends under the threefold axis, leaving the final eleven residues disordered, similarly to the HB1, HB2 and ORF1 only capsids.

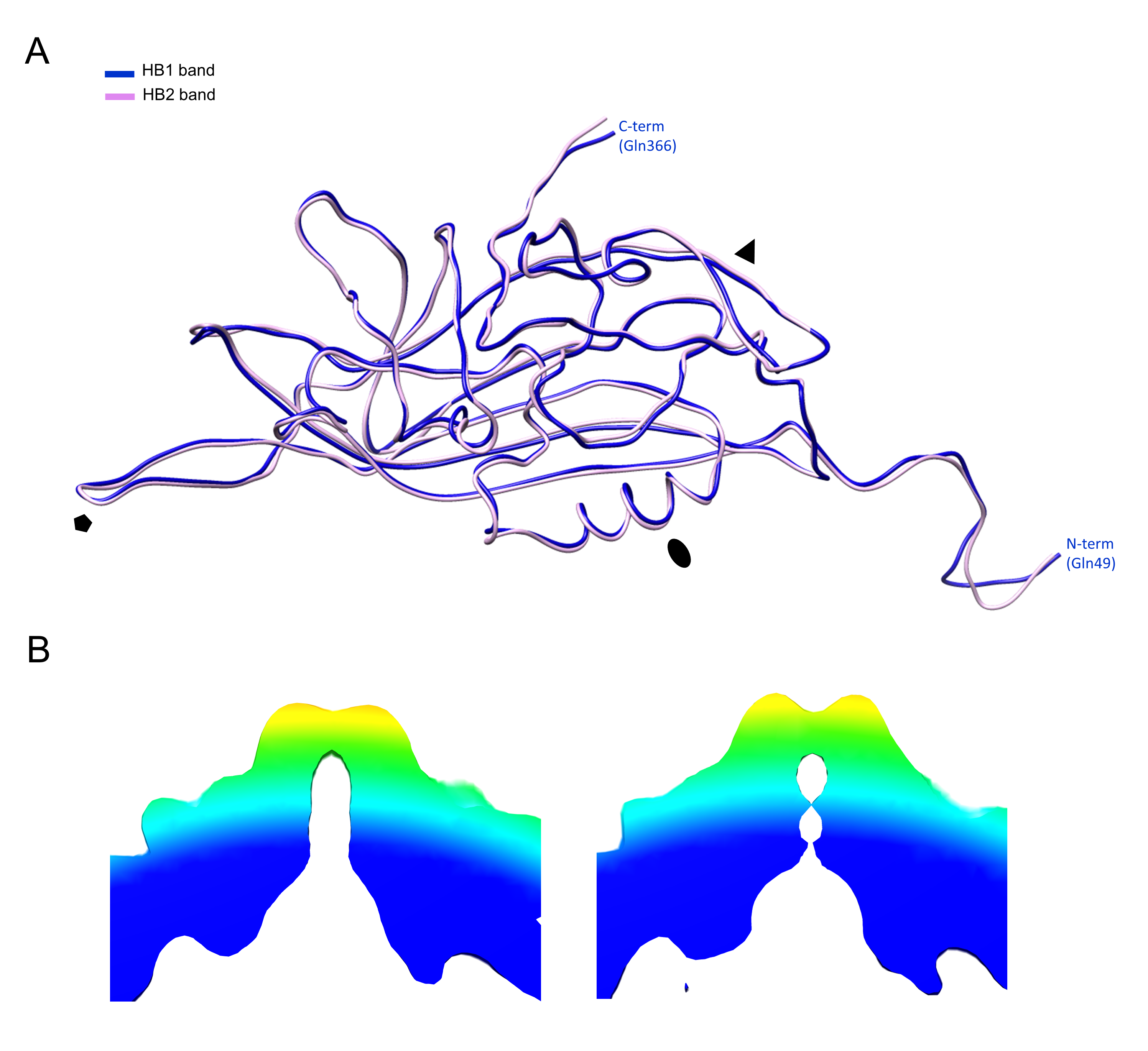

**S9 Figure:** Structural comparison of the two high buoyancy bands (HB1 and HB2, respectively), pulled from the continuous CsCl gradient. (A) Structure of the HB1 and HB2 monomers are identical, with the exception of highly flexible, poorly modelled regions. (B) Low resolution structural reconstruction of the HB1 (left) and HB2 (right) capsids, showing the cross section of the fivefold channel, at the resolution of 8.4 Å and 9.3 Å, respectively (ơ=1). The narrowing of the HB2 fivefold channel could not be observed in the high resolution cryoEM structures.

ORF1 only VLPs

Normalized RFU

| Temperature (◦C) | pH 7.4 | pH 6.0 | pH 5.5 | pH 4.0 |
| --- | --- | --- | --- | --- |
| 30 | 0 | 0.0153838 | 0 | 0 |
| 30.5 | 0 | 0.0333316 | 0 | 0 |
| 31 | 0 | 0.0307677 | 0 | 0 |
| 31.5 | 0 | 0.0307677 | 0 | 0 |
| 32 | 0 | 0.0307677 | 0 | 0 |
| 32.5 | 0 | 0.0322658 | 0 | 0 |
| 33 | 0 | 0.0236364 | 0 | 0 |
| 33.5 | 0 | 0.000651451 | 0 | 0 |
| 34 | 0 | 0 | 0 | 0 |
| 34.5 | 0 | 0 | 0 | 0 |
| 35 | 0 | 0 | 0 | 0 |
| 35.5 | 0 | 0 | 0 | 0 |
| 36 | 0 | 0 | 0 | 0 |
| 36.5 | 0 | 0 | 0 | 0 |
| 37 | 0 | 0 | 0 | 0 |
| 37.5 | 0 | 0 | 0 | 0 |
| 38 | 0 | 0 | 0 | 0 |
| 38.5 | 0 | 0 | 0 | 0 |
| 39 | 0 | 0 | 0 | 0 |
| 39.5 | 0 | 0 | 0 | 0 |
| 40 | 0 | 0 | 0 | 0 |
| 40.5 | 0 | 0 | 0 | 0 |
| 41 | 0 | 0 | 0 | 0 |
| 41.5 | 0 | 0 | 0 | 0 |
| 42 | 0 | 0 | 0 | 0 |
| 42.5 | 0 | 0 | 0 | 0 |
| 43 | 0 | 0 | 0 | 0 |
| 43.5 | 0 | 0 | 0 | 0 |
| 44 | 0.0022319 | 0 | 0 | 0 |
| 44.5 | 0.000661277 | 0 | 0 | 0 |
| 45 | 0.00150257 | 0 | 0 | 0 |
| 45.5 | 0.0125813 | 0 | 0 | 0 |
| 46 | 0.0149365 | 0 | 0 | 0 |
| 46.5 | 0.0153562 | 0 | 0 | 0 |
| 47 | 0.0228811 | 0 | 0 | 0 |
| 47.5 | 0.0244381 | 0 | 0 | 0 |
| 48 | 0.0255116 | 0.000452125 | 0 | 0 |
| 48.5 | 0.031559 | 0.00444135 | 0 | 0 |
| 49 | 0.0321069 | 0.00442122 | 0 | 0 |
| 49.5 | 0.0316744 | 0.0101789 | 0 | 0 |
| 50 | 0.0351136 | 0.00424459 | 0 | 0 |
| 50.5 | 0.032911 | 0 | 0 | 0 |
| 51 | 0.0286558 | 0 | 0 | 0 |
| 51.5 | 0.0254541 | 0 | 0 | 0 |
| 52 | 0.0186501 | 0 | 0 | 0 |
| 52.5 | 0.0135762 | 0 | 0 | 0 |
| 53 | 0.0104854 | 0 | 0.00517301 | 0 |
| 53.5 | 0.0110212 | 0 | 0.00335832 | 0 |
| 54 | 0.016008 | 0 | 0.00291585 | 0 |
| 54.5 | 0.0173495 | 0.00452167 | 0.00917151 | 0 |
| 55 | 0.0117091 | 0.00115549 | 0.0132974 | 0 |
| 55.5 | 0.00402288 | 0 | 0.0124726 | 0 |
| 56 | 0 | 0.00585797 | 0.0152382 | 0 |
| 56.5 | 0 | 0.00667897 | 0.0189264 | 0 |
| 57 | 0 | 0.00169766 | 0.0211164 | 0 |
| 57.5 | 0 | 0 | 0.0199121 | 0 |
| 58 | 0 | 0 | 0.0176031 | 0 |
| 58.5 | 0 | 0 | 0.0164095 | 0 |
| 59 | 0 | 0 | 0.0129316 | 0 |
| 59.5 | 0 | 0 | 0.0108103 | 0 |
| 60 | 0 | 0 | 0.0111215 | 0 |
| 60.5 | 0 | 0 | 0.00883751 | 0 |
| 61 | 0 | 0 | 0.00177279 | 0 |
| 61.5 | 0 | 0 | 0.00426312 | 0 |
| 62 | 0 | 0 | 0.00463153 | 0 |
| 62.5 | 0 | 0 | 0 | 0 |
| 63 | 0 | 0 | 0 | 0 |
| 63.5 | 0 | 0 | 2.89774e-05 | 0 |
| 64 | 0 | 0 | 0 | 0 |
| 64.5 | 0 | 0 | 0 | 0 |
| 65 | 0 | 0 | 0 | 0 |
| 65.5 | 0 | 0 | 0.00683422 | 0 |
| 66 | 0 | 0 | 0.0125428 | 0 |
| 66.5 | 0 | 0 | 0.0149576 | 0 |
| 67 | 0 | 0 | 0.0172092 | 0 |
| 67.5 | 0 | 0 | 0.0140036 | 0 |
| 68 | 0 | 0 | 0.00856292 | 0 |
| 68.5 | 0 | 0 | 0.00217767 | 0 |
| 69 | 0 | 0 | 0 | 0 |
| 69.5 | 0.00435248 | 0 | 0 | 0 |
| 70 | 0.0170877 | 0 | 0 | 0.0460214 |
| 70.5 | 0.0337854 | 0 | 0 | 0.234918 |
| 71 | 0.054584 | 0 | 0 | 0.466415 |
| 71.5 | 0.0933471 | 0 | 0.0110591 | 0.705018 |
| 72 | 0.154095 | 0.00805722 | 0.0216085 | 0.90249 |
| 72.5 | 0.230024 | 0.0237451 | 0.0264268 | 1 |
| 73 | 0.326109 | 0.0365033 | 0.0377389 | 0.946465 |
| 73.5 | 0.442762 | 0.0427348 | 0.0458858 | 0.75036 |
| 74 | 0.571991 | 0.0516694 | 0.0480466 | 0.462988 |
| 74.5 | 0.70918 | 0.0653864 | 0.055165 | 0.133417 |
| 75 | 0.843468 | 0.0853635 | 0.0715989 | 0 |
| 75.5 | 0.946355 | 0.123139 | 0.0950466 | 0 |
| 76 | 1 | 0.188078 | 0.134074 | 0 |
| 76.5 | 0.995608 | 0.285146 | 0.198695 | 0 |
| 77 | 0.913327 | 0.420571 | 0.292181 | 0 |
| 77.5 | 0.759612 | 0.586711 | 0.422842 | 0 |
| 78 | 0.572056 | 0.763631 | 0.586596 | 0 |
| 78.5 | 0.36669 | 0.915837 | 0.76097 | 0 |
| 79 | 0.163408 | 1 | 0.910317 | 0 |
| 79.5 | 0.00752757 | 0.989888 | 1 | 0 |
| 80 | 0 | 0.879753 | 0.99596 | 0 |
| 80.5 | 0 | 0.689389 | 0.882984 | 0 |
| 81 | 0 | 0.458248 | 0.68256 | 0 |
| 81.5 | 0 | 0.236694 | 0.447623 | 0 |
| 82 | 0 | 0.0565786 | 0.217683 | 0 |
| 82.5 | 0 | 0 | 0.0230054 | 0 |
| 83 | 0 | 0 | 0 | 0 |
| 83.5 | 0 | 0 | 0 | 0 |
| 84 | 0 | 0 | 0 | 0 |
| 84.5 | 0 | 0 | 0 | 0 |
| 85 | 0 | 0 | 0 | 0 |
| 85.5 | 0 | 0 | 0 | 0 |
| 86 | 0 | 0 | 0 | 0 |
| 86.5 | 0 | 0 | 0 | 0 |
| 87 | 0 | 0 | 0 | 0 |
| 87.5 | 0 | 0 | 0 | 0 |
| 88 | 0 | 0 | 0 | 0 |
| 88.5 | 0 | 0 | 0 | 0 |
| 89 | 0 | 0 | 0 | 0 |
| 89.5 | 0 | 0 | 0 | 0 |
| 90 | 0 | 0 | 0 | 0 |
| 90.5 | 0 | 0 | 0 | 0 |
| 91 | 0 | 0 | 0 | 0 |
| 91.5 | 0 | 0 | 0 | 0 |
| 92 | 0 | 0 | 0 | 0 |
| 92.5 | 0 | 0 | 0 | 0 |
| 93 | 0 | 0 | 0 | 0 |
| 93.5 | 0 | 0 | 0 | 0 |
| 94 | 0 | 0 | 0 | 0 |
| 94.5 | 0 | 0 | 0 | 0 |
| 95 | 0 | 0 | 0 | 0 |
| 95.5 | 0 | 0 | 0 | 0 |
| 96 | 0 | 0 | 0 | 0 |
| 96.5 | 0 | 0 | 0 | 0 |
| 97 | 0 | 0 | 0 | 0 |
| 97.5 | 0 | 0 | 0 | 0 |
| 98 | 0 | 0 | 0 | 0 |
| 98.5 | 0 | 0 | 0 | 0 |
| 99 | 0 | 0 | 0 | 0 |

High buoyancy band (sucrose)

Normalized RFU

| Temperature (◦C) | pH 7.4 | pH 6.0 | pH 5.5 | pH 4.0 |
| --- | --- | --- | --- | --- |
| 30 | 0 | 0 | 0 | 0.0100272 |
| 30.5 | 0 | 0 | 0 | 0.0217255 |
| 31 | 0 | 0 | 0 | 0.0200544 |
| 31.5 | 0 | 0 | 0 | 0.0200544 |
| 32 | 0 | 0 | 0 | 0.0200544 |
| 32.5 | 0 | 0 | 0 | 0.0202368 |
| 33 | 0 | 0 | 0 | 0.0189611 |
| 33.5 | 0 | 0 | 0 | 0.0175718 |
| 34 | 0 | 0 | 0 | 0.0182695 |
| 34.5 | 0 | 0 | 0 | 0.0198428 |
| 35 | 0 | 0 | 0 | 0.0246859 |
| 35.5 | 0 | 0 | 0 | 0.0284984 |
| 36 | 0 | 0 | 0 | 0.0313542 |
| 36.5 | 0 | 0 | 0 | 0.0328009 |
| 37 | 0 | 0 | 0 | 0.0305735 |
| 37.5 | 0 | 0 | 0 | 0.0285641 |
| 38 | 0 | 0 | 0 | 0.0285421 |
| 38.5 | 0 | 0 | 0 | 0.0288047 |
| 39 | 0 | 0 | 0 | 0.029845 |
| 39.5 | 0 | 0 | 2.28201e-05 | 0.0356678 |
| 40 | 0 | 0 | 0.00753768 | 0.0394509 |
| 40.5 | 0 | 0 | 0.0151422 | 0.0394071 |
| 41 | 0 | 0 | 0.0217011 | 0.0378659 |
| 41.5 | 0 | 0 | 0.0258615 | 0.0370103 |
| 42 | 0 | 0 | 0.0301125 | 0.0332658 |
| 42.5 | 0 | 0 | 0.0351527 | 0.0316017 |
| 43 | 0 | 0 | 0.0362739 | 0.0347135 |
| 43.5 | 0 | 0 | 0.0381761 | 0.0367785 |
| 44 | 0 | 0 | 0.0424213 | 0.0396043 |
| 44.5 | 0 | 0.00213847 | 0.0472919 | 0.0436646 |
| 45 | 0 | 0.00752224 | 0.0521059 | 0.0421657 |
| 45.5 | 0 | 0.0140952 | 0.0606781 | 0.0381846 |
| 46 | 0 | 0.0223678 | 0.0707608 | 0.0379183 |
| 46.5 | 0 | 0.0330181 | 0.0793606 | 0.0328333 |
| 47 | 0.0239891 | 0.0433758 | 0.0853302 | 0.0266566 |
| 47.5 | 0.0550713 | 0.0543762 | 0.0897541 | 0.0257896 |
| 48 | 0.08602 | 0.0634456 | 0.0915188 | 0.0245486 |
| 48.5 | 0.114123 | 0.0718938 | 0.0875377 | 0.0205753 |
| 49 | 0.133283 | 0.0751172 | 0.0823199 | 0.0203363 |
| 49.5 | 0.15035 | 0.0746461 | 0.0775113 | 0.020016 |
| 50 | 0.162039 | 0.0709381 | 0.0717089 | 0.0190438 |
| 50.5 | 0.161024 | 0.0672592 | 0.0643042 | 0.0191357 |
| 51 | 0.155036 | 0.0600414 | 0.0584537 | 0.0171333 |
| 51.5 | 0.147726 | 0.0524058 | 0.053941 | 0.0146326 |
| 52 | 0.133575 | 0.049018 | 0.0486374 | 0.0131119 |
| 52.5 | 0.118559 | 0.047989 | 0.0416543 | 0.0103092 |
| 53 | 0.107221 | 0.0452112 | 0.0360403 | 0.0082213 |
| 53.5 | 0.0939839 | 0.0417659 | 0.0353236 | 0.0104496 |
| 54 | 0.0782577 | 0.0412149 | 0.0348821 | 0.0121575 |
| 54.5 | 0.0660339 | 0.0393544 | 0.0360052 | 0.012027 |
| 55 | 0.0565804 | 0.036251 | 0.041464 | 0.0125782 |
| 55.5 | 0.046164 | 0.0333733 | 0.0462059 | 0.0113019 |
| 56 | 0.0416391 | 0.0324608 | 0.0462887 | 0.00731906 |
| 56.5 | 0.0383959 | 0.0289205 | 0.0464716 | 0.00616103 |
| 57 | 0.0313286 | 0.0248617 | 0.047807 | 0.00752047 |
| 57.5 | 0.0236688 | 0.0227914 | 0.0475528 | 0.00834164 |
| 58 | 0.0200551 | 0.0192308 | 0.048384 | 0.0111023 |
| 58.5 | 0.0134267 | 0.0170601 | 0.0510332 | 0.0150612 |
| 59 | 0.00540543 | 0.0185896 | 0.0533097 | 0.015646 |
| 59.5 | 0.00318375 | 0.0177213 | 0.0538331 | 0.0160853 |
| 60 | 0.00162168 | 0.0158806 | 0.0537797 | 0.0188291 |
| 60.5 | 0 | 0.0198076 | 0.0513757 | 0.0206317 |
| 61 | 0 | 0.0206944 | 0.0467271 | 0.0228448 |
| 61.5 | 0 | 0.0148464 | 0.0402583 | 0.0262502 |
| 62 | 0 | 0.0114641 | 0.0316803 | 0.0290287 |
| 62.5 | 0 | 0.00646163 | 0.0239591 | 0.0316251 |
| 63 | 0 | 0 | 0.0190312 | 0.0343924 |
| 63.5 | 0 | 0 | 0.0145285 | 0.0372759 |
| 64 | 0 | 0 | 0.00976336 | 0.0422425 |
| 64.5 | 0 | 0 | 0.00437525 | 0.0497905 |
| 65 | 0 | 0 | 0 | 0.0568123 |
| 65.5 | 0 | 0 | 0 | 0.0665238 |
| 66 | 0.00056661 | 0 | 0 | 0.0824684 |
| 66.5 | 0 | 0 | 0 | 0.102421 |
| 67 | 0 | 0 | 0 | 0.131345 |
| 67.5 | 0.000423404 | 0 | 0 | 0.183407 |
| 68 | 0 | 0 | 0 | 0.266001 |
| 68.5 | 0.00486607 | 0 | 0 | 0.383645 |
| 69 | 0.0189533 | 0 | 0 | 0.541651 |
| 69.5 | 0.0355185 | 0 | 0 | 0.721938 |
| 70 | 0.0636786 | 0 | 0 | 0.882274 |
| 70.5 | 0.109004 | 0 | 0 | 0.985345 |
| 71 | 0.175737 | 0 | 0.00101361 | 1 |
| 71.5 | 0.266904 | 0 | 0.00895597 | 0.910799 |
| 72 | 0.38341 | 0.00453879 | 0.012272 | 0.734185 |
| 72.5 | 0.524694 | 0.00975242 | 0.0209728 | 0.514837 |
| 73 | 0.681304 | 0.0158325 | 0.026591 | 0.300554 |
| 73.5 | 0.827721 | 0.0311141 | 0.0281689 | 0.125708 |
| 74 | 0.939897 | 0.0494775 | 0.0382967 | 0.00558706 |
| 74.5 | 1 | 0.0821936 | 0.0605438 | 0 |
| 75 | 0.982408 | 0.14806 | 0.093169 | 0 |
| 75.5 | 0.881024 | 0.24406 | 0.151557 | 0 |
| 76 | 0.713546 | 0.37152 | 0.245323 | 0 |
| 76.5 | 0.50592 | 0.538546 | 0.370309 | 0 |
| 77 | 0.288437 | 0.720914 | 0.524866 | 0 |
| 77.5 | 0.0965989 | 0.879332 | 0.701264 | 0 |
| 78 | 0 | 0.984189 | 0.867833 | 0 |
| 78.5 | 0 | 1 | 0.977048 | 0 |
| 79 | 0 | 0.912646 | 1 | 0 |
| 79.5 | 0 | 0.736075 | 0.923759 | 0 |
| 80 | 0 | 0.508146 | 0.753958 | 0 |
| 80.5 | 0 | 0.276561 | 0.524226 | 0 |
| 81 | 0 | 0.0819463 | 0.289707 | 0 |
| 81.5 | 0 | 0 | 0.0947828 | 0 |
| 82 | 0 | 0 | 0 | 0 |
| 82.5 | 0 | 0 | 0 | 0 |
| 83 | 0 | 0 | 0 | 0 |
| 83.5 | 0 | 0 | 0 | 0 |
| 84 | 0 | 0 | 0 | 0 |
| 84.5 | 0 | 0 | 0 | 0 |
| 85 | 0 | 0 | 0 | 0 |
| 85.5 | 0 | 0 | 0 | 0 |
| 86 | 0 | 0 | 0 | 0 |
| 86.5 | 0 | 0 | 0 | 0 |
| 87 | 0 | 0 | 0 | 0 |
| 87.5 | 0 | 0 | 0 | 0 |
| 88 | 0 | 0 | 0 | 0 |
| 88.5 | 0 | 0 | 0 | 0 |
| 89 | 0 | 0 | 0 | 0 |
| 89.5 | 0 | 0 | 0 | 0 |
| 90 | 0 | 0 | 0 | 0 |
| 90.5 | 0 | 0 | 0 | 0 |
| 91 | 0 | 0 | 0 | 0 |
| 91.5 | 0 | 0 | 0 | 0 |
| 92 | 0 | 0 | 0 | 0 |
| 92.5 | 0 | 0 | 0 | 0 |
| 93 | 0 | 0 | 0 | 0 |
| 93.5 | 0 | 0 | 0 | 0 |
| 94 | 0 | 0 | 0 | 0 |
| 94.5 | 0 | 0 | 0 | 0 |
| 95 | 0 | 0 | 0 | 0 |
| 95.5 | 0 | 0 | 0 | 0 |
| 96 | 0 | 0 | 0 | 0 |
| 96.5 | 0 | 0 | 0 | 0 |
| 97 | 0 | 0 | 0 | 0 |
| 97.5 | 0 | 0 | 0 | 0 |
| 98 | 0 | 0 | 0 | 0 |
| 98.5 | 0 | 0 | 0 | 0 |
| 99 | 0 | 0.030336 | 0 | 0 |

High buoyancy band (CsCl) 1

Normalized RFU

| Temperature (◦C) | pH7.4 | pH 6.0 | pH 5.5 | pH 4.0 |
| --- | --- | --- | --- | --- |
| 30 | 0 | 0 | 0.00681409 | 0.0376618 |
| 30.5 | 0 | 0 | 0.0147639 | 0.0816006 |
| 31 | 0 | 0 | 0.0136282 | 0.0753237 |
| 31.5 | 0 | 0 | 0.0136282 | 0.0753237 |
| 32 | 0 | 0 | 0.0136282 | 0.0753237 |
| 32.5 | 0 | 0 | 0.0153246 | 0.0770342 |
| 33 | 0 | 0 | 0.00556269 | 0.065801 |
| 33.5 | 0 | 0 | 0 | 0.0488272 |
| 34 | 0 | 0 | 0 | 0.0438822 |
| 34.5 | 0 | 0 | 0 | 0.0440468 |
| 35 | 0 | 0 | 0 | 0.0486233 |
| 35.5 | 0 | 0 | 0 | 0.0526129 |
| 36 | 0 | 0 | 0 | 0.0512424 |
| 36.5 | 0 | 0 | 0 | 0.046051 |
| 37 | 0 | 0 | 0 | 0.0418715 |
| 37.5 | 0 | 0 | 0 | 0.0396048 |
| 38 | 0 | 0 | 0 | 0.0380181 |
| 38.5 | 0 | 0 | 0 | 0.0382718 |
| 39 | 0 | 0 | 0 | 0.0438265 |
| 39.5 | 0 | 0 | 0 | 0.0458008 |
| 40 | 0 | 0 | 0 | 0.0393171 |
| 40.5 | 0 | 0 | 0 | 0.0348417 |
| 41 | 0 | 0 | 0 | 0.0328147 |
| 41.5 | 0 | 0 | 0 | 0.023598 |
| 42 | 0 | 0 | 0 | 0.0187129 |
| 42.5 | 0 | 0 | 0 | 0.0210189 |
| 43 | 0 | 0 | 0 | 0.0229227 |
| 43.5 | 0 | 0 | 0 | 0.0260587 |
| 44 | 0 | 0 | 0 | 0.0306857 |
| 44.5 | 0 | 0 | 0 | 0.0314686 |
| 45 | 0 | 0 | 0 | 0.0288712 |
| 45.5 | 0 | 0 | 0 | 0.024845 |
| 46 | 0 | 0 | 0 | 0.0227449 |
| 46.5 | 0.00806297 | 0 | 0 | 0.0219569 |
| 47 | 0.0138441 | 0 | 0 | 0.0192919 |
| 47.5 | 0.0209686 | 0 | 0 | 0.018662 |
| 48 | 0.0361721 | 0 | 0 | 0.0184385 |
| 48.5 | 0.052064 | 0 | 0 | 0.0153782 |
| 49 | 0.0607548 | 0 | 0 | 0.013521 |
| 49.5 | 0.0535763 | 0 | 0 | 0.0164149 |
| 50 | 0.0513059 | 0 | 0 | 0.0161373 |
| 50.5 | 0.0461157 | 0 | 0 | 0.0116784 |
| 51 | 0.0306484 | 0 | 0 | 0.00834677 |
| 51.5 | 0.0213715 | 0 | 0 | 0.00519359 |
| 52 | 0.030344 | 0 | 0 | 0 |
| 52.5 | 0.0264417 | 0 | 0 | 0 |
| 53 | 0.0190052 | 0 | 0 | 0 |
| 53.5 | 0.0223339 | 0 | 0 | 0 |
| 54 | 0.0198049 | 0.0233508 | 0 | 0 |
| 54.5 | 0.0143993 | 0.00570992 | 0 | 0.00257483 |
| 55 | 0.0191162 | 0.00699384 | 0 | 0.0115791 |
| 55.5 | 0.0252067 | 0 | 0 | 0.0141284 |
| 56 | 0.022429 | 0 | 0 | 0.0221306 |
| 56.5 | 0.0170446 | 0 | 0 | 0.0292737 |
| 57 | 0.0116722 | 0 | 0 | 0.0256633 |
| 57.5 | 0.00632994 | 0 | 0 | 0.0237187 |
| 58 | 0 | 0 | 0 | 0.0273599 |
| 58.5 | 0.00422119 | 0 | 0 | 0.0262497 |
| 59 | 0.0133825 | 0 | 0 | 0.0251263 |
| 59.5 | 0.0140078 | 0 | 0 | 0.0354462 |
| 60 | 0.0139038 | 0 | 0 | 0.044295 |
| 60.5 | 0.0186031 | 0 | 0 | 0.0474531 |
| 61 | 0.00989352 | 0 | 0 | 0.0557333 |
| 61.5 | 0.00300431 | 0 | 0 | 0.0708032 |
| 62 | 0.00645025 | 0.0301288 | 0 | 0.0775451 |
| 62.5 | 0.00499787 | 0.0838106 | 0 | 0.0853838 |
| 63 | 0.0117556 | 0.124642 | 0 | 0.105934 |
| 63.5 | 0.0356681 | 0.153019 | 0 | 0.120898 |
| 64 | 0.0652684 | 0.159687 | 0 | 0.125688 |
| 64.5 | 0.101018 | 0.134579 | 0 | 0.134088 |
| 65 | 0.141922 | 0.111739 | 0.00661171 | 0.141039 |
| 65.5 | 0.175538 | 0.0826813 | 0.00297432 | 0.134203 |
| 66 | 0.194435 | 0.065671 | 0.00237651 | 0.127846 |
| 66.5 | 0.19461 | 0.0637921 | 0.0028644 | 0.127264 |
| 67 | 0.179655 | 0.0765738 | 0.00810869 | 0.133364 |
| 67.5 | 0.149427 | 0.0970589 | 0.0183578 | 0.158911 |
| 68 | 0.116913 | 0.147207 | 0.0311432 | 0.216691 |
| 68.5 | 0.0955476 | 0.201803 | 0.0548315 | 0.317245 |
| 69 | 0.0794498 | 0.227208 | 0.0987497 | 0.465385 |
| 69.5 | 0.0745377 | 0.232639 | 0.132181 | 0.651554 |
| 70 | 0.102806 | 0.238475 | 0.144605 | 0.832218 |
| 70.5 | 0.134605 | 0.207179 | 0.162764 | 0.961498 |
| 71 | 0.166321 | 0.157292 | 0.161602 | 1 |
| 71.5 | 0.237585 | 0.12806 | 0.132774 | 0.923564 |
| 72 | 0.334654 | 0.128072 | 0.102023 | 0.735611 |
| 72.5 | 0.441317 | 0.129708 | 0.0894743 | 0.490713 |
| 73 | 0.583069 | 0.128823 | 0.0762624 | 0.249104 |
| 73.5 | 0.751581 | 0.14544 | 0.0708539 | 0.0507323 |
| 74 | 0.888622 | 0.176852 | 0.0824843 | 0 |
| 74.5 | 0.976732 | 0.196757 | 0.110954 | 0 |
| 75 | 1 | 0.220952 | 0.146219 | 0 |
| 75.5 | 0.934364 | 0.311069 | 0.202309 | 0 |
| 76 | 0.779226 | 0.44349 | 0.310058 | 0 |
| 76.5 | 0.570461 | 0.59921 | 0.45851 | 0 |
| 77 | 0.350598 | 0.784018 | 0.627739 | 0 |
| 77.5 | 0.152258 | 0.941064 | 0.802313 | 0 |
| 78 | 0 | 1 | 0.9468 | 0 |
| 78.5 | 0 | 0.960756 | 1 | 0 |
| 79 | 0 | 0.825899 | 0.943486 | 0 |
| 79.5 | 0 | 0.61079 | 0.790915 | 0 |
| 80 | 0 | 0.365914 | 0.567301 | 0 |
| 80.5 | 0 | 0.155511 | 0.321248 | 0 |
| 81 | 0 | 0 | 0.108829 | 0 |
| 81.5 | 0 | 0 | 0 | 0 |
| 82 | 0 | 0 | 0 | 0 |
| 82.5 | 0 | 0 | 0 | 0 |
| 83 | 0 | 0 | 0 | 0 |
| 83.5 | 0 | 0 | 0 | 0 |
| 84 | 0 | 0 | 0 | 0 |
| 84.5 | 0 | 0 | 0 | 0 |
| 85 | 0 | 0 | 0 | 0 |
| 85.5 | 0 | 0 | 0 | 0 |
| 86 | 0 | 0 | 0 | 0 |
| 86.5 | 0 | 0 | 0 | 0 |
| 87 | 0 | 0 | 0 | 0 |
| 87.5 | 0 | 0 | 0 | 0 |
| 88 | 0 | 0 | 0 | 0 |
| 88.5 | 0 | 0 | 0 | 0 |
| 89 | 0 | 0 | 0 | 0 |
| 89.5 | 0 | 0 | 0 | 0 |
| 90 | 0 | 0 | 0 | 0 |
| 90.5 | 0 | 0 | 0 | 0 |
| 91 | 0 | 0 | 0 | 0 |
| 91.5 | 0 | 0 | 0 | 0 |
| 92 | 0 | 0 | 0 | 0 |
| 92.5 | 0 | 0 | 0 | 0 |
| 93 | 0 | 0 | 0 | 0 |
| 93.5 | 0 | 0 | 0 | 0 |
| 94 | 0 | 0 | 0 | 0 |
| 94.5 | 0 | 0 | 0 | 0 |
| 95 | 0 | 0 | 0 | 0 |
| 95.5 | 0 | 0 | 0 | 0 |
| 96 | 0 | 0 | 0 | 0 |
| 96.5 | 0 | 0 | 0 | 0 |
| 97 | 0 | 0 | 0 | 0 |
| 97.5 | 0 | 0 | 0 | 0 |
| 98 | 0 | 0 | 0 | 0 |
| 98.5 | 0 | 0.00254535 | 0 | 0 |
| 99 | 0 | 0.0146767 | 0 | 0 |

High buoyancy band (CsCl) 2

Normalized RFU

| Temperature (◦C) | pH7.4 | pH 6.0 | pH 5.5 | pH 4.0 |
| --- | --- | --- | --- | --- |
| 30 | 0 | 0 | 0 | 0.0134602 |
| 30.5 | 0 | 0 | 0 | 0.0291638 |
| 31 | 0 | 0 | 0 | 0.0269204 |
| 31.5 | 0 | 0 | 0 | 0.0269204 |
| 32 | 0 | 0 | 0 | 0.0269204 |
| 32.5 | 0 | 0 | 0 | 0.0286879 |
| 33 | 0 | 0 | 0 | 0.0168844 |
| 33.5 | 0 | 0 | 0 | 0.00186224 |
| 34 | 0 | 0 | 0 | 0 |
| 34.5 | 0 | 0 | 0 | 0 |
| 35 | 0 | 0 | 0 | 0 |
| 35.5 | 0 | 0 | 0 | 0 |
| 36 | 0 | 0 | 0 | 0 |
| 36.5 | 0 | 0 | 0 | 0 |
| 37 | 0 | 0 | 0 | 0.0011965 |
| 37.5 | 0 | 0 | 0 | 0 |
| 38 | 0 | 0 | 0 | 0 |
| 38.5 | 0 | 0 | 0 | 0 |
| 39 | 0 | 0 | 0 | 0 |
| 39.5 | 0 | 0 | 0 | 0 |
| 40 | 0 | 0 | 0 | 0.00743113 |
| 40.5 | 0 | 0 | 0 | 0.0130378 |
| 41 | 0 | 0 | 0 | 0.0188009 |
| 41.5 | 0 | 0 | 0 | 0.0218744 |
| 42 | 0 | 0 | 0 | 0.0144827 |
| 42.5 | 0 | 0 | 0 | 0.00240773 |
| 43 | 0 | 0 | 0 | 0 |
| 43.5 | 0 | 0 | 0 | 0 |
| 44 | 0 | 0 | 0 | 7.60797e-05 |
| 44.5 | 0 | 0 | 0 | 0.0105295 |
| 45 | 0 | 0 | 0 | 0.0212887 |
| 45.5 | 0 | 0 | 0 | 0.0301272 |
| 46 | 0 | 0 | 0 | 0.0320085 |
| 46.5 | 0 | 0 | 0 | 0.0258298 |
| 47 | 0 | 0 | 0 | 0.0181071 |
| 47.5 | 0 | 0 | 0 | 0.0129926 |
| 48 | 0 | 0 | 0 | 0.00966701 |
| 48.5 | 0.00405378 | 0 | 0 | 0.00749087 |
| 49 | 0.0179664 | 0 | 0 | 0.00689428 |
| 49.5 | 0.0212122 | 0 | 0 | 0.00883977 |
| 50 | 0.0212096 | 0 | 0 | 0.0119969 |
| 50.5 | 0.0250088 | 0.00111931 | 0 | 0.00539337 |
| 51 | 0.0257865 | 0.000467196 | 0 | 0 |
| 51.5 | 0.0212938 | 0.00033042 | 0 | 0 |
| 52 | 0.0168317 | 0 | 0 | 0 |
| 52.5 | 0.0218053 | 0 | 0 | 0 |
| 53 | 0.0254464 | 0 | 0 | 0 |
| 53.5 | 0.0260213 | 0 | 0 | 0.00722548 |
| 54 | 0.0281168 | 0 | 0 | 0.0189276 |
| 54.5 | 0.0291643 | 0.000117385 | 0 | 0.0236832 |
| 55 | 0.0218666 | 0.00380951 | 0 | 0.0214944 |
| 55.5 | 0.0164084 | 0 | 0 | 0.0313922 |
| 56 | 0.0138285 | 0 | 0 | 0.0358349 |
| 56.5 | 0.00828953 | 0.00701682 | 0 | 0.0330624 |
| 57 | 0.00922324 | 0.0111933 | 0 | 0.0374399 |
| 57.5 | 0.0150622 | 0.0150769 | 0.00462616 | 0.0444815 |
| 58 | 0.0194385 | 0.0257852 | 0 | 0.042295 |
| 58.5 | 0.0254433 | 0.0303697 | 0 | 0.0406314 |
| 59 | 0.0358922 | 0.0293117 | 0 | 0.0405491 |
| 59.5 | 0.0406359 | 0.0318587 | 0.00904175 | 0.0361916 |
| 60 | 0.0408323 | 0.0379773 | 0.0176624 | 0.0382471 |
| 60.5 | 0.0458689 | 0.0441331 | 0.0277701 | 0.0449057 |
| 61 | 0.0510727 | 0.0463207 | 0.0421474 | 0.0532624 |
| 61.5 | 0.0582427 | 0.0458468 | 0.0427783 | 0.0635354 |
| 62 | 0.070247 | 0.0445808 | 0.0262281 | 0.0767734 |
| 62.5 | 0.0874294 | 0.0450906 | 0.0206672 | 0.0880361 |
| 63 | 0.104433 | 0.042393 | 0.0263355 | 0.0938255 |
| 63.5 | 0.129973 | 0.0428987 | 0.0188046 | 0.0961071 |
| 64 | 0.16062 | 0.0518942 | 0.020203 | 0.10013 |
| 64.5 | 0.192829 | 0.0630037 | 0.0340617 | 0.0987697 |
| 65 | 0.233224 | 0.0666312 | 0.0357623 | 0.0927775 |
| 65.5 | 0.27709 | 0.0743438 | 0.0272392 | 0.0969286 |
| 66 | 0.306344 | 0.0876538 | 0.0423101 | 0.108468 |
| 66.5 | 0.321573 | 0.0988692 | 0.0525918 | 0.11431 |
| 67 | 0.322938 | 0.114261 | 0.0498228 | 0.132528 |
| 67.5 | 0.299995 | 0.145223 | 0.0567551 | 0.170332 |
| 68 | 0.253216 | 0.185476 | 0.0676147 | 0.219535 |
| 68.5 | 0.203389 | 0.224072 | 0.0707031 | 0.308085 |
| 69 | 0.159964 | 0.260179 | 0.081393 | 0.455476 |
| 69.5 | 0.129679 | 0.281338 | 0.106064 | 0.632469 |
| 70 | 0.114799 | 0.28023 | 0.126015 | 0.805629 |
| 70.5 | 0.123908 | 0.253713 | 0.142096 | 0.947924 |
| 71 | 0.161403 | 0.215801 | 0.147075 | 1 |
| 71.5 | 0.221107 | 0.174793 | 0.141229 | 0.92703 |
| 72 | 0.306006 | 0.137077 | 0.124296 | 0.755154 |
| 72.5 | 0.422449 | 0.107309 | 0.0994796 | 0.530535 |
| 73 | 0.574237 | 0.0954004 | 0.0870179 | 0.299701 |
| 73.5 | 0.730252 | 0.0995271 | 0.0873065 | 0.104911 |
| 74 | 0.869593 | 0.114344 | 0.0835085 | 0 |
| 74.5 | 0.968676 | 0.149058 | 0.092828 | 0 |
| 75 | 1 | 0.215731 | 0.138645 | 0 |
| 75.5 | 0.933338 | 0.311378 | 0.193382 | 0 |
| 76 | 0.786806 | 0.437781 | 0.274402 | 0 |
| 76.5 | 0.583949 | 0.601305 | 0.421021 | 0 |
| 77 | 0.35309 | 0.778718 | 0.598096 | 0 |
| 77.5 | 0.139892 | 0.922893 | 0.770425 | 0 |
| 78 | 0 | 1 | 0.920344 | 0 |
| 78.5 | 0 | 0.983758 | 1 | 0 |
| 79 | 0 | 0.85405 | 0.962689 | 0 |
| 79.5 | 0 | 0.637207 | 0.821138 | 0 |
| 80 | 0 | 0.385297 | 0.605359 | 0 |
| 80.5 | 0 | 0.149376 | 0.36222 | 0 |
| 81 | 0 | 0 | 0.144149 | 0 |
| 81.5 | 0 | 0 | 0 | 0 |
| 82 | 0 | 0 | 0 | 0 |
| 82.5 | 0 | 0 | 0 | 0 |
| 83 | 0 | 0 | 0 | 0 |
| 83.5 | 0 | 0 | 0 | 0 |
| 84 | 0 | 0 | 0 | 0 |
| 84.5 | 0 | 0 | 0 | 0 |
| 85 | 0 | 0 | 0 | 0 |
| 85.5 | 0 | 0 | 0 | 0 |
| 86 | 0 | 0 | 0 | 0 |
| 86.5 | 0 | 0 | 0 | 0 |
| 87 | 0 | 0 | 0 | 0 |
| 87.5 | 0 | 0 | 0 | 0 |
| 88 | 0 | 0 | 0 | 0 |
| 88.5 | 0 | 0 | 0 | 0 |
| 89 | 0 | 0 | 0 | 0 |
| 89.5 | 0 | 0 | 0 | 0 |
| 90 | 0 | 0 | 0 | 0 |
| 90.5 | 0 | 0 | 0 | 0 |
| 91 | 0 | 0 | 0 | 0 |
| 91.5 | 0 | 0 | 0 | 0 |
| 92 | 0 | 0 | 0 | 0 |
| 92.5 | 0 | 0 | 0 | 0 |
| 93 | 0 | 0 | 0 | 0 |
| 93.5 | 0 | 0 | 0 | 0 |
| 94 | 0 | 0 | 0 | 0 |
| 94.5 | 0 | 0 | 0 | 0 |
| 95 | 0 | 0 | 0 | 0 |
| 95.5 | 0 | 0 | 0 | 0 |
| 96 | 0 | 0 | 0 | 0 |
| 96.5 | 0 | 0 | 0 | 0 |
| 97 | 0 | 0 | 0 | 0 |
| 97.5 | 0 | 0 | 0 | 0 |
| 98 | 0 | 0 | 0 | 0 |
| 98.5 | 0 | 0 | 0 | 0 |
| 99 | 0 | 0 | 0 | 0 |

Low buoyancy band (sucrose)

Normalized RFU

| Temperature (◦C) | pH 7.4 | pH 6.0 | pH 5.5 | pH 4.0 |
| --- | --- | --- | --- | --- |
| 30 | 0 | 0 | 0 | 0 |
| 30.5 | 0 | 0 | 0 | 0 |
| 31 | 0 | 0 | 0 | 0 |
| 31.5 | 0 | 0 | 0 | 0 |
| 32 | 0 | 0 | 0 | 0 |
| 32.5 | 0 | 0 | 0 | 0 |
| 33 | 0 | 0 | 0 | 0 |
| 33.5 | 0 | 0 | 0 | 0 |
| 34 | 0 | 0 | 0 | 0 |
| 34.5 | 0 | 0 | 0 | 0 |
| 35 | 0 | 0 | 0 | 0.00308866 |
| 35.5 | 0 | 0 | 0 | 0.0241912 |
| 36 | 0 | 0 | 0 | 0.0369657 |
| 36.5 | 0 | 0 | 0 | 0.0357569 |
| 37 | 0 | 0 | 0 | 0.0301387 |
| 37.5 | 0 | 0 | 0 | 0.0235718 |
| 38 | 0 | 0 | 0 | 0.0171012 |
| 38.5 | 0 | 0 | 0 | 0.00920109 |
| 39 | 0 | 0 | 0 | 0.0118896 |
| 39.5 | 0 | 0 | 0 | 0.0238445 |
| 40 | 0 | 0 | 0 | 0.0248667 |
| 40.5 | 0 | 0 | 0 | 0.0197451 |
| 41 | 0 | 0 | 0 | 0.0226911 |
| 41.5 | 0 | 0 | 0 | 0.0203196 |
| 42 | 0 | 0 | 0 | 0.0136894 |
| 42.5 | 0 | 0 | 0 | 0.017986 |
| 43 | 0 | 0 | 0 | 0.025855 |
| 43.5 | 0 | 0 | 0 | 0.028032 |
| 44 | 0 | 0 | 0 | 0.027248 |
| 44.5 | 0 | 0 | 0 | 0.0243448 |
| 45 | 0 | 0 | 0 | 0.01936 |
| 45.5 | 0 | 0 | 0 | 0.0143465 |
| 46 | 0 | 0 | 0 | 0.0125859 |
| 46.5 | 0 | 0 | 0 | 0.0215742 |
| 47 | 0 | 0.00612241 | 0 | 0.0285853 |
| 47.5 | 0 | 0.0255927 | 0 | 0.0281406 |
| 48 | 0 | 0.0279599 | 0 | 0.0292789 |
| 48.5 | 0 | 0.0139595 | 0 | 0.029848 |
| 49 | 0 | 0 | 0 | 0.0211321 |
| 49.5 | 0 | 0 | 0 | 0.0246359 |
| 50 | 0 | 0 | 0 | 0.0391253 |
| 50.5 | 0 | 0 | 0 | 0.0480867 |
| 51 | 0 | 0 | 0 | 0.058277 |
| 51.5 | 0 | 0 | 0 | 0.0716468 |
| 52 | 0 | 0 | 0 | 0.0662162 |
| 52.5 | 0 | 0.00239095 | 0 | 0.0516934 |
| 53 | 0 | 0.023131 | 0.00330642 | 0.0435428 |
| 53.5 | 0 | 0.0243053 | 9.8068e-05 | 0.0284613 |
| 54 | 0 | 0.0200577 | 0.00262679 | 0.0123721 |
| 54.5 | 0 | 0.019106 | 0 | 0.00778372 |
| 55 | 0 | 0.00390186 | 0 | 0.00308511 |
| 55.5 | 0 | 0 | 0 | 0 |
| 56 | 0 | 0 | 0 | 0.00480868 |
| 56.5 | 0 | 0 | 0.00590933 | 0.0109785 |
| 57 | 0 | 0 | 0.00313288 | 0.0177297 |
| 57.5 | 0 | 0 | 0.00424603 | 0.0237572 |
| 58 | 0 | 0.0051606 | 0.0129763 | 0.0221701 |
| 58.5 | 0 | 0.00739544 | 0.00645272 | 0.0146305 |
| 59 | 0 | 0.0163394 | 0 | 0.00385163 |
| 59.5 | 0 | 0.0246459 | 0 | 0 |
| 60 | 0 | 0.0351234 | 0.0110731 | 0 |
| 60.5 | 0 | 0.0452561 | 0.0254019 | 0 |
| 61 | 0 | 0.0565696 | 0.0237016 | 0 |
| 61.5 | 0 | 0.0553182 | 0.0311607 | 0.00669449 |
| 62 | 0 | 0.046475 | 0.0467563 | 0.012184 |
| 62.5 | 0 | 0.0352108 | 0.0478596 | 0.00566923 |
| 63 | 0 | 0.0301478 | 0.0305697 | 0 |
| 63.5 | 0 | 0.0401026 | 0.0294783 | 0 |
| 64 | 0 | 0.0595179 | 0.0461142 | 0 |
| 64.5 | 0 | 0.0699676 | 0.0559092 | 0 |
| 65 | 0 | 0.0688489 | 0.0490938 | 0 |
| 65.5 | 0 | 0.0677611 | 0.0492791 | 0.00810636 |
| 66 | 0 | 0.0483902 | 0.0658164 | 0.0131084 |
| 66.5 | 0 | 0.0302076 | 0.0521667 | 0.00560938 |
| 67 | 0 | 0.0392417 | 0.0355707 | 0.00735596 |
| 67.5 | 0 | 0.0557378 | 0.0472083 | 0.0143715 |
| 68 | 0 | 0.0579916 | 0.0643493 | 0.0233474 |
| 68.5 | 0.0142895 | 0.0565192 | 0.0580362 | 0.0343714 |
| 69 | 0.0313025 | 0.0537988 | 0.0508866 | 0.050075 |
| 69.5 | 0.0503724 | 0.0406687 | 0.0460611 | 0.0638809 |
| 70 | 0.0527731 | 0.0218718 | 0.0363345 | 0.0772984 |
| 70.5 | 0.04569 | 0.0193619 | 0.0151837 | 0.0974225 |
| 71 | 0.0508089 | 0.0355581 | 0.00115708 | 0.125298 |
| 71.5 | 0.064769 | 0.0410564 | 0.0146801 | 0.162999 |
| 72 | 0.0996702 | 0.0339863 | 0.0221538 | 0.218177 |
| 72.5 | 0.162536 | 0.0349821 | 0.00823623 | 0.30439 |
| 73 | 0.2408 | 0.0249621 | 0.00402179 | 0.42031 |
| 73.5 | 0.312702 | 0.00636067 | 0.00748449 | 0.571709 |
| 74 | 0.390986 | 0.00230651 | 0 | 0.745835 |
| 74.5 | 0.474364 | 0.00289727 | 0 | 0.900956 |
| 75 | 0.56328 | 0.0107593 | 0.00997891 | 0.993516 |
| 75.5 | 0.664786 | 0.0335652 | 0.0363946 | 1 |
| 76 | 0.779832 | 0.0541557 | 0.0551923 | 0.905182 |
| 76.5 | 0.891731 | 0.0694053 | 0.0814295 | 0.718436 |
| 77 | 0.970728 | 0.101001 | 0.102615 | 0.496639 |
| 77.5 | 1 | 0.130937 | 0.125926 | 0.284541 |
| 78 | 0.957544 | 0.154031 | 0.160181 | 0.109074 |
| 78.5 | 0.849087 | 0.186428 | 0.18747 | 0 |
| 79 | 0.690554 | 0.235626 | 0.20197 | 0 |
| 79.5 | 0.505115 | 0.294245 | 0.238691 | 0 |
| 80 | 0.314826 | 0.359112 | 0.28138 | 0 |
| 80.5 | 0.152904 | 0.443836 | 0.312511 | 0 |
| 81 | 0.026664 | 0.553282 | 0.388264 | 0 |
| 81.5 | 0 | 0.670463 | 0.511083 | 0 |
| 82 | 0 | 0.784094 | 0.632633 | 0 |
| 82.5 | 0 | 0.899376 | 0.756524 | 0 |
| 83 | 0 | 0.979588 | 0.883763 | 0 |
| 83.5 | 0 | 1 | 0.967094 | 0 |
| 84 | 0 | 0.949997 | 1 | 0 |
| 84.5 | 0 | 0.82515 | 0.96715 | 0 |
| 85 | 0 | 0.629548 | 0.857363 | 0 |
| 85.5 | 0 | 0.410446 | 0.695393 | 0 |
| 86 | 0 | 0.191564 | 0.493254 | 0 |
| 86.5 | 0 | 0.0115783 | 0.268708 | 0 |
| 87 | 0 | 0 | 0.0672722 | 0 |
| 87.5 | 0 | 0 | 0 | 0 |
| 88 | 0 | 0 | 0 | 0 |
| 88.5 | 0 | 0 | 0 | 0 |
| 89 | 0 | 0 | 0 | 0 |
| 89.5 | 0 | 0 | 0 | 0 |
| 90 | 0 | 0 | 0 | 0 |
| 90.5 | 0 | 0 | 0 | 0 |
| 91 | 0 | 0 | 0 | 0 |
| 91.5 | 0 | 0 | 0 | 0 |
| 92 | 0 | 0 | 0 | 0 |
| 92.5 | 0 | 0 | 0 | 0 |
| 93 | 0 | 0 | 0 | 0 |
| 93.5 | 0 | 0 | 0 | 0 |
| 94 | 0 | 0 | 0 | 0 |
| 94.5 | 0 | 0 | 0 | 0 |
| 95 | 0 | 0 | 0 | 0 |
| 95.5 | 0 | 0 | 0 | 0 |
| 96 | 0 | 0 | 0 | 0 |
| 96.5 | 0 | 0 | 0 | 0 |
| 97 | 0 | 0 | 0 | 0 |
| 97.5 | 0 | 0 | 0 | 0 |
| 98 | 0 | 0 | 0 | 0 |
| 98.5 | 0 | 0 | 0 | 0 |
| 99 | 0 | 0 | 0 | 0 |

Low buoyancy band (CsCl)

Normalized RFU

| Temperature (◦C) | pH7.4 | pH 6.0 | pH 5.5 | pH 4.0 |
| --- | --- | --- | --- | --- |
| 30 | 0 | 0.0116435 | 0.000682945 | 0 |
| 30.5 | 0 | 0.0252275 | 0.00147972 | 0 |
| 31 | 0 | 0.023287 | 0.00136589 | 0 |
| 31.5 | 0 | 0.023287 | 0.00136589 | 0 |
| 32 | 0 | 0.023287 | 0.00136589 | 0 |
| 32.5 | 0 | 0.025254 | 0.0031773 | 0 |
| 33 | 0 | 0.0130546 | 0 | 0 |
| 33.5 | 0 | 0 | 0 | 0 |
| 34 | 0 | 0 | 0 | 0 |
| 34.5 | 0 | 0 | 0 | 0 |
| 35 | 0 | 0 | 0 | 0.00308866 |
| 35.5 | 0 | 0 | 0 | 0.0241912 |
| 36 | 0 | 0 | 0 | 0.0369657 |
| 36.5 | 0 | 0 | 0 | 0.0357569 |
| 37 | 0 | 0 | 0 | 0.0301387 |
| 37.5 | 0 | 0 | 0 | 0.0235718 |
| 38 | 0 | 0 | 0 | 0.0171012 |
| 38.5 | 0 | 0 | 0 | 0.00920109 |
| 39 | 0 | 0 | 0 | 0.0118896 |
| 39.5 | 0 | 0 | 0 | 0.0238445 |
| 40 | 0 | 0 | 0 | 0.0248667 |
| 40.5 | 0 | 0 | 0 | 0.0197451 |
| 41 | 0 | 0 | 0 | 0.0226911 |
| 41.5 | 0 | 0 | 0 | 0.0203196 |
| 42 | 0 | 0 | 0 | 0.0136894 |
| 42.5 | 0 | 0 | 0 | 0.017986 |
| 43 | 0 | 0 | 0 | 0.025855 |
| 43.5 | 0 | 0 | 0 | 0.028032 |
| 44 | 0 | 0 | 0 | 0.027248 |
| 44.5 | 0 | 0 | 0 | 0.0243448 |
| 45 | 0 | 0 | 0 | 0.01936 |
| 45.5 | 0 | 0 | 0 | 0.0143465 |
| 46 | 0 | 0 | 0 | 0.0125859 |
| 46.5 | 0 | 0 | 0 | 0.0215742 |
| 47 | 0 | 0 | 0 | 0.0285853 |
| 47.5 | 0 | 0 | 0 | 0.0281406 |
| 48 | 0 | 0 | 0 | 0.0292789 |
| 48.5 | 0 | 0 | 0 | 0.029848 |
| 49 | 0 | 0 | 0 | 0.0211321 |
| 49.5 | 0 | 0 | 0 | 0.0246359 |
| 50 | 0 | 0 | 0 | 0.0391253 |
| 50.5 | 0 | 0 | 0 | 0.0480867 |
| 51 | 0 | 0 | 0 | 0.058277 |
| 51.5 | 0 | 0 | 0 | 0.0716468 |
| 52 | 0 | 0 | 0 | 0.0662162 |
| 52.5 | 0 | 0 | 0 | 0.0516934 |
| 53 | 0 | 0 | 0 | 0.0435428 |
| 53.5 | 0 | 0.00605252 | 0 | 0.0284613 |
| 54 | 0 | 0.00608743 | 0 | 0.0123721 |
| 54.5 | 0 | 0 | 0 | 0.00778372 |
| 55 | 0 | 0 | 0 | 0.00308511 |
| 55.5 | 0 | 0 | 0 | 0 |
| 56 | 0 | 0 | 0 | 0.00480868 |
| 56.5 | 0 | 0 | 0 | 0.0109785 |
| 57 | 0 | 0 | 0 | 0.0177297 |
| 57.5 | 0 | 0 | 0 | 0.0237572 |
| 58 | 0 | 0.00460694 | 0 | 0.0221701 |
| 58.5 | 0 | 0.0130239 | 0 | 0.0146305 |
| 59 | 0 | 0.00354142 | 0 | 0.00385163 |
| 59.5 | 0 | 0 | 0 | 0 |
| 60 | 0 | 0 | 0.00457065 | 0 |
| 60.5 | 0 | 0 | 0.00472545 | 0 |
| 61 | 0 | 0 | 0 | 0 |
| 61.5 | 0 | 0.00120898 | 0.00909889 | 0.00669449 |
| 62 | 0 | 0.0158336 | 0.0142488 | 0.012184 |
| 62.5 | 0 | 0.0246488 | 0.00476826 | 0.00566923 |
| 63 | 0 | 0.0221965 | 0.0118446 | 0 |
| 63.5 | 0 | 0.00836435 | 0.0261097 | 0 |
| 64 | 0 | 0.0022128 | 0.0185544 | 0 |
| 64.5 | 0 | 0.00872273 | 0.0217441 | 0 |
| 65 | 0 | 0.00638285 | 0.0285277 | 0 |
| 65.5 | 0 | 0.019896 | 0.0207904 | 0.00810636 |
| 66 | 0.0565222 | 0.0451059 | 0.011205 | 0.0131084 |
| 66.5 | 0.0684502 | 0.0538512 | 0.00527931 | 0.00560938 |
| 67 | 0.0697023 | 0.0492902 | 0.00052365 | 0.00735596 |
| 67.5 | 0.0647163 | 0.0547691 | 0.00854264 | 0.0143715 |
| 68 | 0.0276369 | 0.0562077 | 0.0202966 | 0.0233474 |
| 68.5 | 0 | 0.0465826 | 0.0213188 | 0.0343714 |
| 69 | 0 | 0.0431198 | 0.0261182 | 0.050075 |
| 69.5 | 0 | 0.0443128 | 0.0292708 | 0.0638809 |
| 70 | 0 | 0.0466682 | 0.0202034 | 0.0772984 |
| 70.5 | 0 | 0.0585694 | 0.019563 | 0.0974225 |
| 71 | 0 | 0.0735329 | 0.0413909 | 0.125298 |
| 71.5 | 0.0901104 | 0.0861361 | 0.0647925 | 0.162999 |
| 72 | 0.201835 | 0.105545 | 0.0804955 | 0.218177 |
| 72.5 | 0.319959 | 0.125713 | 0.101286 | 0.30439 |
| 73 | 0.443913 | 0.128321 | 0.112895 | 0.42031 |
| 73.5 | 0.572576 | 0.126232 | 0.118817 | 0.571709 |
| 74 | 0.691571 | 0.122128 | 0.132044 | 0.745835 |
| 74.5 | 0.804001 | 0.113917 | 0.145563 | 0.900956 |
| 75 | 0.911205 | 0.101327 | 0.167954 | 0.993516 |
| 75.5 | 0.974892 | 0.0925922 | 0.190099 | 1 |
| 76 | 1 | 0.103543 | 0.188872 | 0.905182 |
| 76.5 | 0.947972 | 0.122956 | 0.18041 | 0.718436 |
| 77 | 0.798686 | 0.140177 | 0.183949 | 0.496639 |
| 77.5 | 0.588838 | 0.164616 | 0.169645 | 0.284541 |
| 78 | 0.335953 | 0.20361 | 0.172075 | 0.109074 |
| 78.5 | 0.0281432 | 0.254053 | 0.209385 | 0 |
| 79 | 0 | 0.321641 | 0.252579 | 0 |
| 79.5 | 0 | 0.401751 | 0.300357 | 0 |
| 80 | 0 | 0.505524 | 0.367357 | 0 |
| 80.5 | 0 | 0.629994 | 0.43984 | 0 |
| 81 | 0 | 0.741689 | 0.514025 | 0 |
| 81.5 | 0 | 0.843893 | 0.610339 | 0 |
| 82 | 0 | 0.946808 | 0.716865 | 0 |
| 82.5 | 0 | 1 | 0.823727 | 0 |
| 83 | 0 | 0.970846 | 0.919485 | 0 |
| 83.5 | 0 | 0.872129 | 0.989109 | 0 |
| 84 | 0 | 0.706382 | 1 | 0 |
| 84.5 | 0 | 0.483985 | 0.94792 | 0 |
| 85 | 0 | 0.239343 | 0.834619 | 0 |
| 85.5 | 0 | 0.0243557 | 0.66057 | 0 |
| 86 | 0 | 0 | 0.450693 | 0 |
| 86.5 | 0 | 0 | 0.229305 | 0 |
| 87 | 0 | 0 | 0.0359652 | 0 |
| 87.5 | 0 | 0 | 0 | 0 |
| 88 | 0 | 0 | 0 | 0 |
| 88.5 | 0 | 0 | 0 | 0 |
| 89 | 0 | 0 | 0 | 0 |
| 89.5 | 0 | 0 | 0 | 0 |
| 90 | 0 | 0 | 0 | 0 |
| 90.5 | 0 | 0 | 0 | 0 |
| 91 | 0 | 0 | 0 | 0 |
| 91.5 | 0 | 0 | 0 | 0 |
| 92 | 0 | 0 | 0 | 0 |
| 92.5 | 0 | 0 | 0 | 0 |
| 93 | 0 | 0 | 0 | 0 |
| 93.5 | 0 | 0 | 0 | 0 |
| 94 | 0 | 0 | 0 | 0 |
| 94.5 | 0 | 0 | 0 | 0 |
| 95 | 0 | 0 | 0 | 0 |
| 95.5 | 0 | 0 | 0 | 0 |
| 96 | 0 | 0 | 0 | 0 |
| 96.5 | 0 | 0 | 0 | 0 |
| 97 | 0 | 0 | 0 | 0 |
| 97.5 | 0 | 0 | 0 | 0 |
| 98 | 0 | 0 | 0 | 0 |
| 98.5 | 0 | 0 | 0 | 0 |
| 99 | 0 | 0 | 0 | 0 |

**S10 Figure:** Chlorometric assay measuring the PLA2 activity of the Acheta domestica segmented densovirus (AdSDV) capsids, using parvovirus B19 virus like particles (VLPs) as positive control. Values are normalized to the ones measured at the first timepoint of the assay. HB abbreviates the high buoyancy capsids pulled from the 20% band of the sucrose step gradient, used for AdSDV purification. LB (low buoyancy) capsids derive from the 30% band of the aforementioned sucrose step gradient. ORF1 indicates VLPs assembled exclusively from the protein product of VP ORF1, lacking the PLA2 domain, and functions as negative control. Purified particles were heated to 60^◦^C to obtain the data indicated as “heated”.
